# Fine-scale landscape genomics helps explain the slow spread of *Wolbachia* through the *Aedes aegypti* population in Cairns, Australia

**DOI:** 10.1101/103598

**Authors:** Thomas. L. Schmidt, Igor. Filipović, Ary A. Hoffmann, Gordana Rašić

## Abstract

The endosymbiotic bacterium *Wolbachia* suppresses the capacity for arboviral transmission in the mosquito *Aedes aegypti*, and can spread through wild mosquito populations following local introductions. Recent introductions in Cairns, Australia have demonstrated slower than expected spread, that could be due to: i) barriers to *Ae. aegypti* dispersal; ii) leptokurtically distributed dispersal distances; and iii) intergenerational loss of *Wolbachia*. We investigated these three potential causes using genome-wide single-nucleotide polymorphisms (SNPs) and an assay for the *Wolbachia* infection *w*Mel in 161 *Ae. aegypti* collected from Cairns in 2015. We observed a significant barrier effect of Cairns highways on *Ae. aegypti* dispersal using distance-based redundancy analysis and patch-based simulation analysis. We detected putative full-siblings in ovitraps 1312m apart, suggesting long-distance female movement likely mediated by human transport. Finally, we found a pair of full-siblings of different infection status, suggesting loss of *Wolbachia* in the field. While the long-distance movement and *Wolbachia* loss currently represent single observations, these findings together with the identified dispersal barriers can contribute to the slow spread of *Wolbachia* through the *Ae. aegypti* population in Cairns. Our landscape genomics approach can be extended to other host/symbiont systems that are being considered for biocontrol.

## Introduction

The mosquito *Aedes aegypti* (Diptera, Culicinae) is the primary vector of arboviruses such as dengue, Zika and chikungunya, that impose an increasing burden on human health worldwide [1, 2], Conventional approaches to combatting these viruses have involved the suppression of *Ae. aegypti* populations through source reduction or insecticide-based programs, but these have had limited efficacy [3, 4], One alternative strategy involves the release of *Ae. aegypti* carrying an infection of the virus-inhibiting, endosymbiotic bacteria *Wolbachia* into wild *Ae. aegypti* populations [5]. Once they are widespread in host populations, *Wolbachia* are expected to diminish the disease transmission rate enough to prevent outbreaks [6]. Although *Ae. aegypti* does not naturally carry *Wolbachia*, several *Wolbachia* strains have been successfully transferred into *Ae. aegypti* from other hosts [7, 8].

The *Wolbachia* strain *w*Mel, originating from *D. melanogaster*, has proven suitable for field deployments in *Ae. aegypti* given its viral blockage [6], moderate fitness costs [7], high intergenerational transmission fidelity [9] and complete cytoplasmic incompatibility [7]. Cytoplasmic incompatibility (CI) describes the phenomenon that offspring of uninfected females mated with *Wolbachia-infected* males are unviable, while offspring of *Wolbachia-infected* females are viable and will carry the infection regardless of male infection status [10]. When *Wolbachia-infected* males are common, CI greatly reduces the relative fitness of uninfected females, which ensures that future generations will be increasingly comprised of *Wolbachia-infected* individuals [11]. However, *Wolbachia* also impose frequency-independent fitness costs on hosts, which for *w*Mel in *Ae. aegypti* include a shorter lifespan [7] and reduced larval competitive ability [9, 12–14], The interaction between costs and benefits produces a critical frequency of *Wolbachia* infection 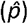 that needs to be exceeded for *Wolbachia* to invade the mosquito population [11,15, 16].

Releases of *w*Mel-infected *Ae. aegypti* into two sites near Cairns, Australia, confirmed that *Wolbachia* can establish stably in quasi-isolated habitat patches [9, 17]. A subsequent study with releases centred within continuous mosquito habitat demonstrated the successful spread of the invasion into surrounding habitat, so that the infection region grew ≈70-85% over the ensuing 18 months [18]. However, the rate of spread of approximately 100-200m per year is close to the lower bound of predicted spread speeds for this host/*Wolbachia* system [19]. This suggests that various potential biological and environmental processes could be operating to restrict the speed or extent of the spread [19, 20].

This study investigates three potential causes for the slow spread observed in Cairns: i) barriers to mosquito dispersal; ii) leptokurtic distribution of dispersal distances; and iii) occasional intergenerational loss of *Wolbachia* (designated as *μ*). Identifying causes of slow spread in Cairns will help future *Wolbachia* deployments to achieve regional invasion faster and at lower cost. We screened *Ae. aegypti* from Cairns for the *w*Mel transinfection and genotyped individuals at genome-wide single nucleotide polymorphisms (SNPs) using double digest restriction-site-associated DNA sequencing (ddRADseq [21]). SNP datasets produced with ddRADseq have been used to elucidate genetic structure in *Ae. aegypti* within cities [22, 23], and have power superior to microsatellites when inferring relationships between *Ae. aegypti* individuals and populations [24],

Fine-scale dispersal of *Ae. aegypti* is mostly accomplished through short-range flight [25–30], which may make highways and other geographical features effective barriers to movement. In Cairns, a Mark-Release-Recapture (MRR) study with releases centred next to a 20m-wide road recorded lower recapture rates at traps across the road [28]. Similarly, patches separated by a 120 m-wide highway in Trinidad had different frequencies of mitochondrial haplotypes [31]. Observing *Wolbachia* invasion dynamics around highways also allows for indirect inference of highway barrier effects. Specifically, *Wolbachia* failed to invade a region across a highway in Gordonvale, Queensland after several years [19], and similar dynamics were recently observed in urban Cairns [18]. In this study we explicitly tested the hypothesis that the genetic structure of *Ae. aegypti* in Cairns is affected by highways acting as barriers to dispersal. This was tested alongside the hypotheses that genetic structure reflected a simple isolation-by-distance (IBD [32]) pattern, and that it was affected by the recent patchy and asynchronous releases of *Wolbachia-infected* mosquitoes throughout the region (see Schmidt et al. [18]).

If host dispersal distances follow a high-kurtosis, leptokurtic distribution, *Wolbachia* spread can proceed up to ≈4 times slower than if the distribution has low kurtosis [19]. This is because high-kurtosis distributions have greater numbers of short-distance and long-distance dispersers [33]. Longer-than-average dispersal distances increase the likelihood that a *Wolbachia-infected* individual will move to an uninvaded region, where they will receive no frequency-dependent fitness benefits and thus contribute less to the invasion. Passive dispersal of *Ae. aegypti* by humans, such as when mosquitoes enter vehicles, may lead to higher kurtosis. Passive movement may often be orders of magnitude higher than the typical 50-100m flight range of *Ae. aegypti* [25–30], leading to higher kurtosis of the dispersal kernel. Passive dispersal is commonly observed in immatures [34], however, movement of imagoes may also occur. *Aedes aegypti* females lay batches of eggs over multiple gonotrophic cycles [35], and through multiple acts of “skip” oviposition within a gonotrophic cycle [36], and thus full-sibling immatures can be found at different sites. In this study we use the distance of separation between sampled full-siblings to indicate female flight ranges or passive dispersal distances within one or two gonotrophic cycles.

Slow *Wolbachia* spread can also result from intergenerational loss of *Wolbachia* (i.e. *μ* > 0), which will lead to an increase in the critical infection frequency 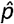 [16]. Loss of *w*Mel has not been detected in laboratory-reared *Ae. aegypti* [7, 13]. However, laboratory populations subjected to high, fluctuating temperatures similar to those of Cairns showed significant decreases in *w*Mel density [37, 38], which could lead to loss of the infection [39, 40]. We looked for evidence of infection loss in Cairns *Ae. aegypti* by first identifying all matrilineages in which the infection was present, then looking for uninfected individuals within these matrilineages. These uninfected individuals will have lost or failed to inherit the infection.

## Methods

### Study site and sample collection

We deployed 110 ovitraps within properties of consenting householders in Cairns, Australia between 13 and 16 April, 2015. Traps covered a 3.3km × 1.9km region of central Cairns, which we partitioned into 6 “plots” for reference: Cairns North West (CNW), Cairns North East (CNE), Parramatta Park North (PPN), Parramatta Park South (PPS), Westcourt (WC) and Bungalow (BN) (Fig 1). Partitions were defined by geographic location, location of highways, and the release history of *w*Mel. Each of these had three possible assignments: the locational groupings of Cairns North [CNW, CNE], Parramatta Park [PPN, PPS] and Westcourt/Bungalow [WC, BN]; the highway groupings of southeast of Bruce Highway [PPS, BN], west of both highways [CNW, PPN, WC], or northeast of Captain Cook Highway [CNE]; and the *Wolbachia* release groupings of releases in 2013 [PPN, WC], releases in 2014 [CNW, BN], or no release history [CNE, PPS] (see Schmidt et al. [18]).

**Fig 1:**
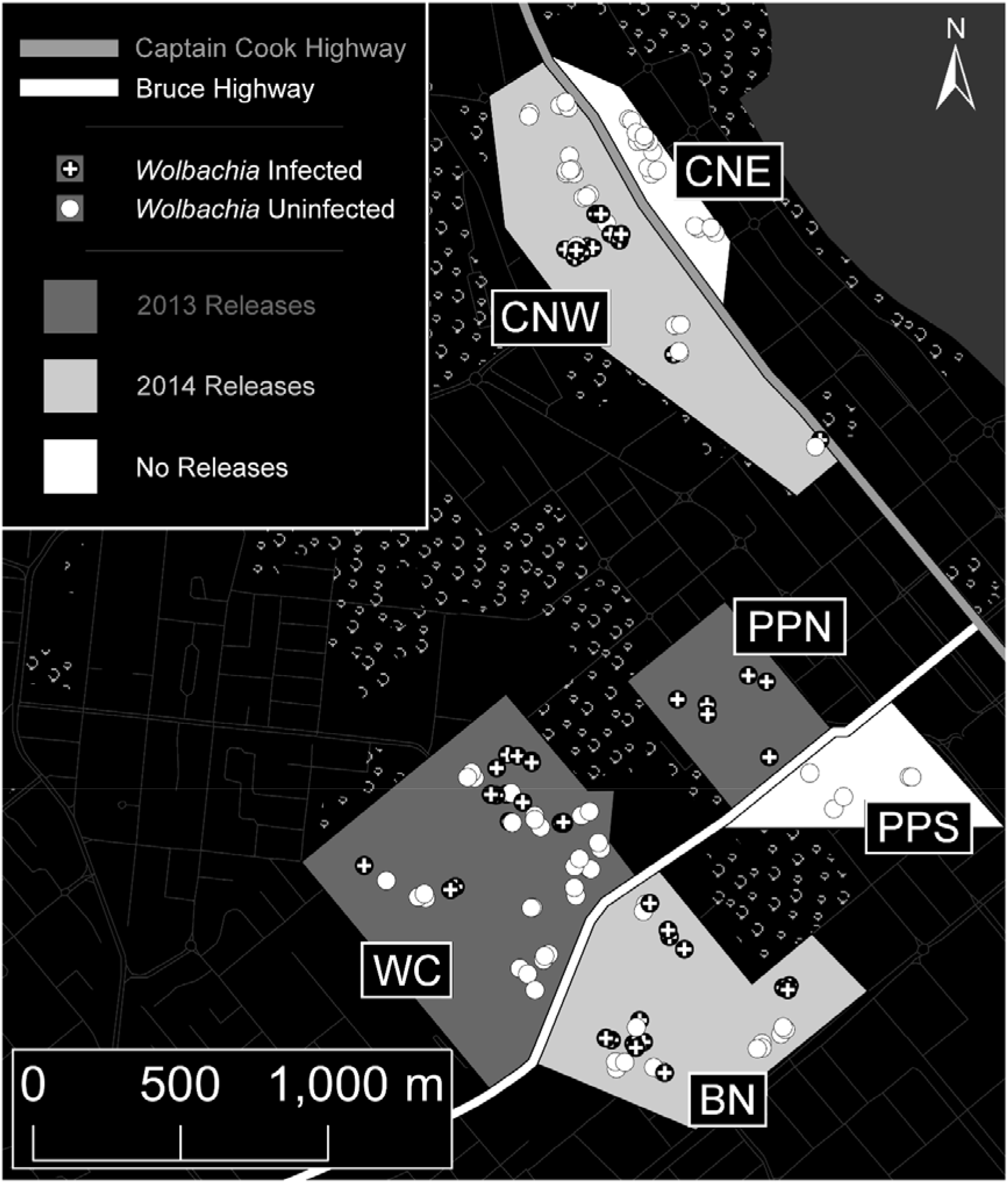
Sampling locations of the mosquitoes analysed with ddRADseq, set within the six sampling plots. Each sample was assigned a *Wolbachia* infection status, a score indicating its position relative to the two highways, and a score indicating when *Wolbachia* releases were carried out in the area. Plot abbreviations are: CNW (Cairns North West), CNE (Cairns North East), PPN (Parramatta Park North), PPS (Parramatta Park South), WC (Westcourt) and BN (Bungalow). (The underlying road network is derived from “Australia Oceania Continent Roads” made available by MapCruzin.com and OpenStreetMap.org under the Open Database License [https://opendatacommons.org/licenses/odbl/1.0/].)

Our sampling period of mid-April was at the end of the region’s monsoonal wet season, when *Ae. aegypti* are abundant [18]. Each ovitrap consisted of a 9.3L black plastic bucket filled halfway with an infusion of water and alfalfa pellets to attract gravid female *Ae. aegypti* [41], which oviposit on strips of red felt clipped to the bucket and extending into the liquid. Traps were left in place for 5-7 days, then the felt strips were removed and dried. Dried strips of mosquito eggs were transferred to the laboratory and hatched by immersion into vessels filled with reverse osmosis water, 2-3 grains of yeast and one quarter of a crushed tablet of tropical fish food. The water, food and yeast were replaced after three days. Emerging virgin imagoes were transferred to freezing ethanol and stored at −20°C until DNA extraction.

We extracted genomic DNA using Roche DNA Isolation Kit for Cells and Tissues (Roche, Pleasanton, CA, USA), with an additional step of RNase treatment. Of the 110 ovitraps deployed, 74 produced *Ae. aegypti* imagoes, from which we selected 161 individuals for sequencing. As we expected ovitraps to contain many full-siblings from the same oviposition [9, 24 42], which can bias analyses of population structure [43], we limited the number of samples per ovitrap to three individuals. This ensured that, after removing closely related individuals, we retained a large enough sample for a powerful analysis of genetic structure.

### ddRADseq library preparation and SNP discovery

We applied the method of Rašić et al. [24] for ddRADseq library preparation, but selected a smaller size range (350-450bp) of genomic fragments to accommodate more individuals per library. Three libraries containing 161 *Ae. aegypti* in total were sequenced in three Illumina HiSeq2500 lanes using 100bp paired-end chemistry.

We processed raw fastq sequences within a customized pipeline [24], retaining reads with phred scores ≥ 13 and trimming them to 90bp. High-quality reads were aligned to the *Ae. aegypti* nuclear genome assembly AaegL1 [44] with the program Bowtie [45]. We allowed for up to three mismatches in the alignment seed, and uniquely aligned reads were analysed using Stacks [46], which we used to call genotypes at RAD stacks of a minimum depth of five reads.

We used the Stacks program *populations* to export VCF files, which were filtered with the program VCFtools [47], We first removed individuals with >20% missing data, leaving 134 individuals. We then removed all loci not in Hardy-Weinberg equilibrium and with minor allele frequencies <0.05. To avoid using markers in high linkage disequilibrium, we applied thinning to ensure no SNP was within 250kbp of another. As *Aedes* genome is thought to contain approximately 2.1 megabases per cM [48], 250kbp roughly corresponds to eight SNPs per map unit, a sampling density that has been shown to largely eradicate the effects of linkage in SNPs [49]. We retained 3,784 unlinked and informative SNPs for analyses of relatedness and genetic structure.

### Assessing barriers to dispersal: Landscape resistance modelling

We tested the hypotheses that highways affect *Ae. aegypti* genetic structure in Cairns (H_1_), and that genetic structure was affected by recent releases of *Wolbachia* (H_2_). For H_1_, each individual mosquito was assigned a score (0, 1, 2) based on its position southeast of Bruce Highway, west of both highways, or northeast of Captain Cook Highway (Fig 1). The difference between a pair of scores represented the number of highways between the two individuals; we called this variable “Highways”. For testing H_2_ each individual was assigned a score (0, 1, 2) based on the *Wolbachia* release history in the area where it was sampled; we called this variable “Releases”. Plots having undergone releases in 2013 (0) were considered more ‘distant’ from those never having undergone releases (2) than those with releases in 2014 (1). We assumed that potential structuring effects such as selection or reductions in diversity would display a greater effect over time. Treating the geographical or temporal separation of individuals as additive variables analogous to landscape resistance surfaces avoids issues associated with detecting discrete barriers [50].

We performed partial distance-based redundancy analyses (dbRDA [51]), to test H_1_ and H_2_ as explicit hypotheses, while controlling for the potentially confounding effects of IBD and latitudinal and longitudinal patterns. We sampled one individual from each matrilineage (see below) with the lowest percentage of missing data, retaining 100 individuals. Easting and Northing UTM coordinates, and a binary variable indicating *Wolbachia* infection status for each individual, called “Infection Status”, were treated as potentially confounding variables and were placed inside a conditional matrix. The dependent variable was a distance matrix of Rousset’s *a* scores [52] calculated for each pair of individuals using the program SPAGeDi [53].

All remaining model procedures were performed in the package VEGAN [54], The dbRDA models were built using the function *capscale*. For all models, we applied the effects of the conditional matrix described above, and assessed the significance of “Highways” and “Releases”. We built three models: one implementing both predictor variables and the other two implementing each variable in isolation. We assessed the marginal significance of each predictor variable with the function *anova.cca*, using 99999 permutations. We then repeated the procedure with “Infection Status” used as a predictor variable alongside “Highways” and “Releases”. While constructing models, we calculated variance-inflation factors (VIF) to check for multicollinearity between predictor variables, using the function *vif.cca*. All VIFs were < 1.1, so none were rejected.

We tested the sensitivity of the variables used to investigate H_1_ and H_2_ by performing additional dbRDAs that followed the above procedure but with modifications made to “Highways” and “Releases”. In each case, instead of a given barrier being 100% the strength of the other, it was assigned strengths of 150% and 200%. Thus, instead of scores of (0, 1, 2) for a given variable, these scores were (0, 1.5, 2.5) or (0, 1, 2.5) for 150% strengths, and (0, 2, 3) or (0, 1, 3) for 200% strengths.

### Assessing barriers to dispersal: Type I error testing

Although ordination methods such as dbRDA can detect genetic structure with greater power than traditional Mantel tests [55, 56], they can also generate more Type I errors [57], Type I errors are common when hypotheses of geographical structure are only proposed against null hypotheses of panmixia, rather than against alternative geographical hypotheses [58, 59]. We considered IBD an alternative hypothesis of genetic structure. A Mantel test performed with the function *mantel* in the R package VEGAN showed a weak correlation between geographical and genetic distances (r = 0.047, P < 0.05), and stronger correlations between geographical distance and both “Highways” (r = 0.424, P < 0.001) and “Releases” (r = 0.305, P < 0.001). When genetic structure exhibits IBD, spatial dependence of barrier variables and inadequate sampling can lead to significance being observed when no barrier effect exists [60]. Therefore, in order to confirm any significance among variables in dbRDA, further analyses were required to eliminate IBD as a hypothesis of genetic structure.

We adapted the method of Kierepka and Latch [60] to evaluate whether the spatial distribution of our traps relative to barriers would lead to a high risk of Type I error. We used CDPOP [61] to simulate the field site without any highways, *Wolbachia* infections, or mosquito release histories, thus creating an artificial environment in which only IBD could explain the pattern of structuring. Parameters and methodology of our CDPOP simulations are detailed in Supplementary Information A and our CDPOP input file is supplied in Supplementary Information B. To summarise: we used the method of Kimura and Crow [62] to calculate the number of effective alleles in our empirical data set, as effective allele counts have been found to correspond well with analytical power across different types of genetic data [63, 64], We assigned genotypes with equivalent allele counts to spatially-referenced individuals corresponding to the 100 individuals used in empirical analyses, and ran simulations for a sufficient number of generations to produce IBD of similar strength to that in the empirical data (r_simulated_, x□ = 0.082, σ = 0.022, all P < 0.05; r_empirical_ = 0.047, P < 0.05). We constructed new dbRDA models using the same parameters as the empirical data, and calculated the marginal significance of “Highways” and “Releases” with ANOVAs. If more than 5 of the 100 simulated samples showed significance for a variable then observations regarding that variable would be at elevated risk of Type I error.

### Long-distance host movement and infection loss: Estimating relatedness/kinship

We used SPAGeDi to calculate Loiselle’s *k* [65] among individuals. First-degree kin relations (full-sibling or parent/offspring) can be ascertained with hundreds of SNPs [66–68], and with the thousands of SNPs available through ddRADseq this confidence can be extended to include second-degree relations (half-sibling or grandparent/grandchild) [24, 69–71], Considering that the 5-7 day sampling period provided by our cross-sectional study design was shorter than the 14 days required for *Ae. aegypti* development [35], we assumed that related individuals were of the same generation. Therefore, we considered pairs with first-degree levels of relatedness to be full-siblings and those with second-degree relatedness to be half-siblings.

We assigned each pair to the most likely kinship category, so that pairs with kinship coefficient *k* > 0.1875 represented putative full-siblings, and those with 0.1875 > *k* > 0.09375 represented putative half-siblings [72], We considered half-siblings to be paternal because polyandry is much rarer than polygyny in wild *Ae. aegypti* [73]. Therefore, we assumed full-siblings to come from the same matrilineage, and half-siblings from different matrilineages.

As our investigations of long-distance movement and *Wolbachia* loss both require precise assignment to kinship categories, we also used the program ML-Relate [74] to run specific hypothesis tests of putative relationships. These were run for all pairs of putative full-siblings of different infection status, and for all individual pairs with *k* > 0.09375 collected from different ovitraps. For each pair we ran one standard test that estimated the relationship assuming that the kinship category assigned using *k* was more likely than the next most likely kinship category, followed by a conservative test that assumed that the kinship category assigned using *k* was less likely to be correct. All tests were run using 10,000,000 simulations.

We considered the distance of separation between full-siblings to indicate movement of adult females. Full-siblings collected from the same ovitrap would likely be from the same gonotrophic cycle, while those collected from different ovitraps could be from one or two gonotrophic cycles. Although passive dispersal cannot be directly inferred from kinship observations, a study near Cairns estimated the average daily movement speed of female *Ae. aegypti* as 17-25m/day [26] and the maximum time between ovitrap deployment and sampling in this study was 7 days. Nevertheless, the distinction between active and passive dispersal is irrelevant to *Wolbachia* invasion dynamics, which depend on the summation of all movement across all life history stages [19, 20]; in this sense, our definition of dispersal is derived from Howard [75]. We thus consider full-sibling separation distances as composites of active flight and passive movement, which are more informative for understanding *Wolbachia* invasion dynamics than active flight alone.

### Long-distance host movement and infection loss: Wolbachia infection screening

Mosquitoes were screened for *Wolbachia* using the protocol of Lee et al. [76]. The infection is diagnosed with PCRs run on the Roche LightCycler^®^ 480 system (384-well format). For each mosquito, PCR was performed using three primer sets, *Aedes* universal primers (*mRpS6*_F/*mRpS6*_R), *Ae. aegypti*-specific primers (*aRpS6*_F/*aRpS6*_R) and *Wolbachia*-specific primers (*wl*_F/*w1*_R). A sample was scored as *Wolbachia* positive when there was robust amplification of all three primer sets *mRpS6, aRpS6* and *w1*, while an *Ae. aegypti* sample that was *Wolbachia* negative amplified only *mRpS6*, and *aRpS6*. Each PCR was run using three positive *Wolbachia* controls and three negative *Wolbachia* controls. *Wolbachia* titre, defined as the ratio of *Wolbachia* gene copies to host gene copies, was estimated using 2^[cp(A)-cp(W)]^, where cp(A) is the crossing point of the *aRpS6* marker and cp(W) is the crossing point of the *w1* marker [76].

We performed three PCR replicates for each individual to confirm its *w*Mel infection status. No inconsistencies in assigning *Wolbachia* infection status were observed across replicates of samples and controls, and similar titres were also observed across replicates (single factor ANOVA *F*-value = 0.341, P = 0.711). Considering the relative consistency of titres across replicates, we used the mean of the three scores to attain final estimates of *Wolbachia* titre for each individual.

From our *Wolbachia*-infection screening and relatedness analysis, we identified *w*Mel-infected matrilineages as being any group of full-siblings containing at least one individual infected with *Wolbachia*. An uninfected individual within an infected matrilineage was considered to have either failed to inherit the infection from its mother or to have lost it during development. We estimated the rate of infection loss per generation *(μ)* by comparing the number of these individuals with the total number of full-siblings from infected matrilineages.

## Results

### Barriers to dispersal

ANOVAs performed on partial dbRDA models showed that only the “Highways” variable was predictive of genetic structure (Table 1; 1.55 ≤ F-value ≤ 1.64; P < 0.05), and was significant irrespective of whether other variables were included. Based on partial Eta squared (η_P_^2^ [77]), “Highways” explained 1.7% of the variance in each model. Of the 100 CDPOP simulations, only two showed significant structuring by “Highways” at the P < 0.05 level, and inclusion of the “Releases” variable in the models did not change this observation. This suggests that the effect of “Highways” observed in the real data is unlikely a Type I error caused by sampling bias or residual autocorrelation. Sensitivity analyses demonstrated robust results for “Highways”, with η_P_^2^ effect sizes in the range 1.54-1.90%.

**Table 1:** Results of ANOVAs testing marginal significance of “Highways” and “Releases” variables in dbRDA. The two variables were each analysed in isolation in separate models, then together in a single model. In every case, “Highways” was predictive of genetic structure while “Releases” was not. Partial Eta squared (η_p_^2^) showed that “Highways” accounted for 1.7% of the variation within each model.

| | | sum of squares | F-value | P | $\eta_p^2$ |
| --- | --- | --- | --- | --- | --- |
| analysed in isolation | HIGHWAYS | 0.021 | 1.635 | **0.021** | 0.017 |
|  | RELEASES | 0.016 | 1.241 | 0.163 | 0.013 |
| analysed together | HIGHWAYS | 0.020 | 1.555 | **0.032** | 0.017 |
|  | RELEASES | 0.015 | 1.165 | 0.231 | 0.012 |

Although η_P_^2^ = 0.017 suggests only a small structuring effect, *Wolbachia* invasions with bistable dynamics are particularly sensitive to barrier effects [20]. Sensitivity increases exponentially as 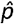 approaches 0.5, and the 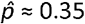 of *w*Mel in *Ae. aegypti* [19] indicates such an invasion can be stopped by a barrier half the strength of that required to stop an invasion with 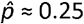 [20]. Likewise, higher 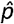 means that slight disruptions to dispersal can slow an advancing wave considerably [19, 20]. To demonstrate this, we used a patch-based simulator derived from Nemo2 [78] and empirical data from Schmidt et al. [18] to simulate the spread of *w*Mel in the absence of barriers. Parameters, methodology and results are detailed in Supplementary Information C. Our simulations showed that, when no barrier effect was set, simulations overestimated infection frequencies in patches separated by highways relative to empirical infection frequencies recorded in Schmidt et al. [18], with an average overestimation across treatments of 6.5%. In comparison, no overestimation of infection frequencies was observed in patches not separated by highways. Overall, simulation of the *Wolbachia* invasion progress south of Bruce Highway (Supplementary Information C) indicated that barrier strength corresponded to an added 30-35 m of separation.

### Estimates of long-distance movement

We detected 31 putative full-sibling groups that contained 43 full-sibling pairs, 41 of which were found within single ovitraps. Two pairs were spread between different plots (Fig 2): one was split between CNW and CNE (239 m separation, *k* = 0.203); the other between CNW and PPS (1312 m separation, *k* = 0.235). Standard maximum-likelihood simulations gave strong support to both pairs being full-siblings (both P < 0.0001). However, conservative simulations were unable to reject the hypotheses that either pair represented half-siblings (P < 0.0001 and P < 0.02 respectively), but rejected them as unrelated (both P = 1).

**Fig 2:**
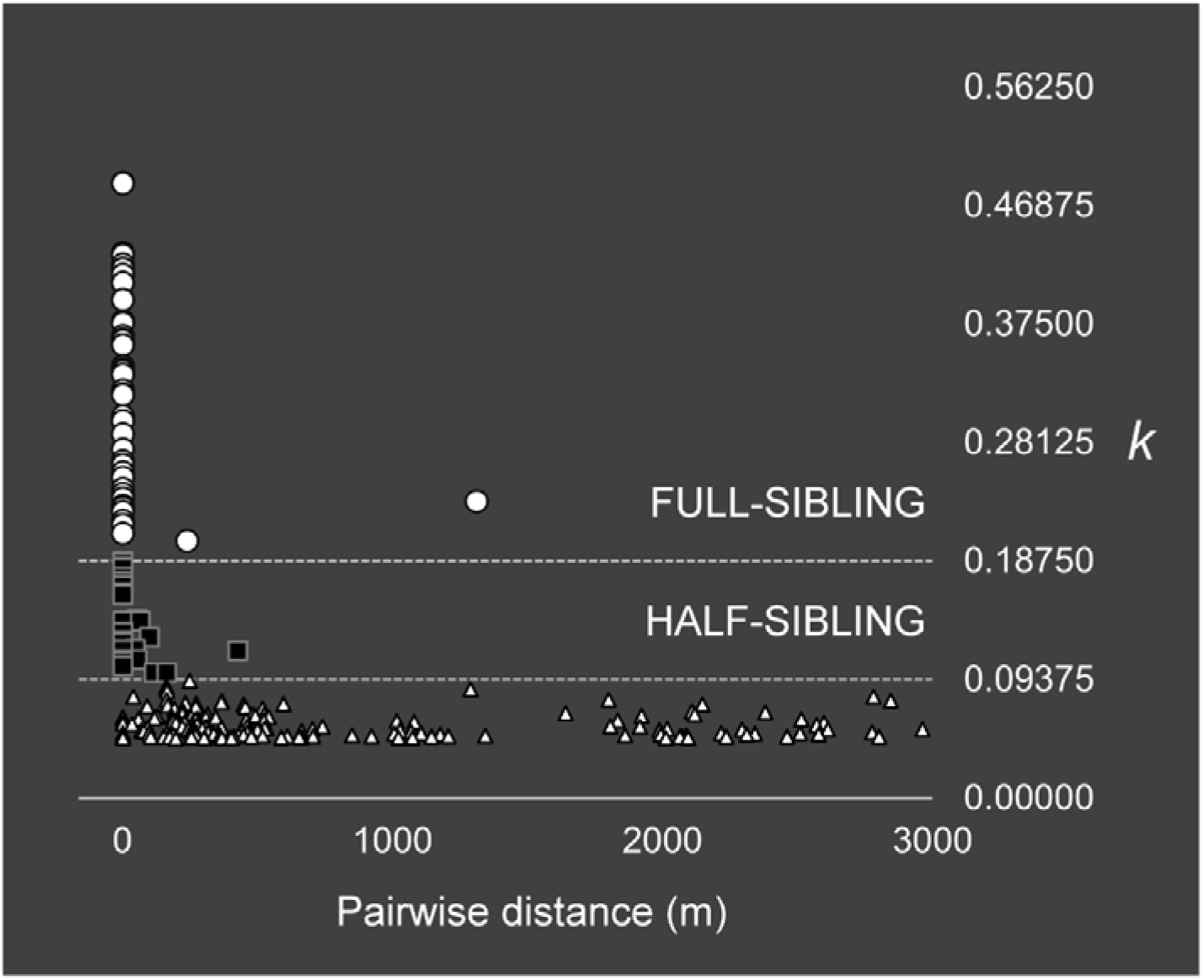
Loiselle’s *k* estimates for sample pairs of relatedness *k* > 0.046875. Pairs of 0.09375 < *k* < 0.1875 are most likely half-sibs, those of *k* < 0.1875 are most likely full-sibs. Most related pairs were found within the same trap, but separation distances of up to 1312 m were observed.

Eight of the 27 putative half-sibling pairs were found across multiple traps (47 - 560 m separation) but not across different plots. Of the eight pairs, only the most closely related pair (*k* = 0.1412, 54m separation, P < 0.07) was confidently determined to be half-siblings and not unrelated using conservative maximum likelihood estimation.

### Loss of Wolbachia

Triplicate PCR runs for *w*Mel detection indicated that 60 of the 161 mosquitoes were infected with *Wolbachia*. The six individuals from PPN were all infected, none of PPS and CNE were infected, and the remaining plots had a mixture of infection rates (Fig 1). We detected 10 *Wolbachia-infected* matrilineages, containing 21 individuals. Importantly, we recorded a single case of infection loss in CNW. The sample from this matrilineage consisted of a pair of putative full-siblings (*k* = 0.376), one of which carried the infection (titre = 6.15) and one of which did not. Maximum likelihood simulation confirmed that this pair represented full-siblings (P < 0.0001) and rejected the alternative hypotheses (both P = 1). We calculated a tentative probability of infection loss among offspring within infected matrilineages, giving a likelihood of loss of one in 21 (*μ* = 0.048), although the 95% binomial confidence intervals around this estimate were large (0.001, 0.238).

Supplementary Information D describes a comparison between *Wolbachia* titres in the lab-reared Cairns sample and that of a field-collected sample from the neighbouring town of Gordonvale. The field-collected sample had titres that were almost three times higher than Cairns on average, but also three times as dispersed (x□ = 18.4, σ = 10.8; compared with x□ = 6.6, σ = 3.8 in Cairns), with one field-reared individual having a titre of only 1 *Wolbachia* gene copy for every 3 *Aedes* gene copies. The lowest titre in the Cairns samples was more than 3 times this amount. The highest titre in Gordonvale was 56 *Wolbachia* gene copies for each *Aedes* gene copy, more than double the highest recorded in Cairns.

## Discussion

Our study produced three main findings: i) highways exert a significant influence on *Ae. aegypti* genetic structure; ii) gravid *Ae. aegypti* females can travel > 1 km; and (iii) *Ae. aegypti* in Cairns may occasionally lose the *w*Mel infection 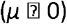. Each phenomenon is expected to have a slowing effect on *w*Mel invasion [19, 20]. Because ii) and iii) were based on single observations, we cannot confidently predict their frequency among *Ae. aegypti* in Cairns. However, field observations of slow spread of *w*Mel through Cairns [18, 19] are congruent with some long-distance movement and occasional infection loss.

### Highways act as dispersal barriers

While previous studies suggested that roads could be barriers to *Ae. aegypti* dispersal [28, 31], we provide the first evidence for such an effect through explicit hypothesis testing within a landscape genetics framework. We detected a small but statistically significant barrier effect of highways, corresponding to 1.7% of dbRDA variance in genetic distance between individuals. Our simulations of *Wolbachia* invasion progress south of Bruce Highway (Supplementary Information C) showed that barrier strength corresponded to an added 30-35m of separation. The *w*Mel invasion observed at PP was slow (100-200m per year [18]), with the infection frequencies at the wave front being only slightly above the critical frequency 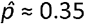 [19]. Therefore, even a small added “cost” to cross the highway could slow the invasion considerably [20]. If the restrictive effect of highways on dispersal increase with highway width and traffic levels, then many urban highways in cities earmarked for future *w*Mel releases would likely be effective barriers to spread.

On the other hand, the restrictive effects of highways on dispersal can strengthen the invasion within the area they enclose. The habitat patches along these highway boundaries will have increased infection frequencies relative to patches in regions that are not subdivided, and could fortify the local invasion from the counteracting effect of uninfected immigrants. Following the 2013 *Wolbachia* releases in Cairns, the largest and most successfully invaded release site at Edge Hill/Whitfield recorded an influx of uninfected *Ae. aegypti* at the start of the 2014/2015 wet season [18]. This site was deemed particularly vulnerable to such reinvasion, due to its greater connectivity to surrounding uninfected regions and very low dry-season mosquito density. Ideally, future release sites in areas with low dry-season densities should be positioned adjacent to dispersal barriers such as highways, as this may help reduce the threat of reinvasion.

### Long-distance movement of Aedes aegypti adult females

Analyses of local kinship patterns have provided several useful inferences regarding *Ae. aegypti* ecology. The pair of probable full-siblings collected from ovitraps 1312m apart indicates movement of a single female between oviposition events, and represents either the extreme end of the flight range in *Ae. aegypti* or some combination of active and passive dispersal. Previous maximum dispersal estimates of gravid *Ae. aegypti* females have been less than a kilometre in a single gonotrophic (egg producing) cycle [79, 80]. In our study, 7 days passed between deployment and sampling of the two ovitraps 1312m apart, which is long enough for an additional unobserved egg deposition somewhere between them [35]. If we assume this distance to reflect active dispersal, an additional gonotrophic cycle is likely. The length of a single such cycle is rarely more than a few days, particularly when temperatures exceed 30°C [35] as they did during our sampling. If the distance was crossed in a single gonotrophic cycle, the average daily speed would have been an order of magnitude greater than the previous estimates of average female flight speed near Cairns of 17-25m/day [26]. Therefore, passive, human-assisted transport is likely to have occurred. Adults of a related mosquito *Aedes albopictus*, have been found in cars stopped at major roads in Barcelona, Spain (David Roiz, pers. comm.)

Using a continuous coefficient of relationship such as *k* to assign pairs of individuals to discrete kinship categories can be problematic when scores are close to the critical cut-off values. Pairing this method with conservative maximum likelihood estimation resolved some of the clearer distinctions (i.e. the full-siblings exhibiting *Wolbachia* loss) but not others (i.e. the putative full-siblings exhibiting long-distance movement). However, the *k* scores of putative full-siblings and putative half-siblings were clearly separated from each other relative to the variability in *k* scores within each category. This reflects the power of genome-wide SNPs for inferring relationships [66–68]. Inferring dispersal from relatedness also avoids potential biases resulting from lab-raised *Ae. aegypti* used in MRR studies failing to develop experience in local conditions that will inform their future oviposition choices [81, 82], Our findings are broadly consistent with the results of several MRR studies showing potential for long distance movement in *Ae. aegypti* imagoes [79, 80], and suggest that this method provides an alternative to MRR for studying dispersal.

### Infection loss in Cairns Aedes aegypti

We found the first evidence in support of intergenerational infection loss (*μ* = 4.8%) in the *Ae. aegypti/w*Mel system, albeit from a single data point. It is worth noting that a comparable loss of 3.3% was recorded for the *w*AlbA infection in field-collected *Ae. albopictus* [83]. Transmission of *w*Mel in *Ae. aegypti* has previously been estimated as perfect or quasi-perfect in the field, but this was based on assaying eggs that were oviposited in the laboratory [9]. By comparison, samples in this study were eclosed from eggs that had spent days in the field before collection, potentially exposing them to stressors such as high heat fluctuations, which have been observed to affect titre of *w*Mel transmission in laboratory populations [37], Fluctuating high temperatures could be the reason for the very low titres recorded for some Gordonvale individuals (Supplementary Information D), though this could also be due a reduction in titre with age, which has been recorded for *w*AlbA in *Ae. albopictus* males [84], The inclusion of blood-fed females in the Gordonvale sample may explain the occurrence of some very high *w*Mel titres, which are known to double in blood-fed *Ae. aegypti* [85].

The *w*Mel strain successfully invaded *Ae. aegypti* populations in Gordonvale and Yorkey’s Knob, two quasi-isolated release sites near Cairns, in 2011 [17] and these areas have maintained infection rates close to 95% for years without reaching fixation [9], Under the assumptions of *μ* = 0, high migration rates of uninfected gravid females would be required to maintain the observed frequencies of infection of 0.03 into Gordonvale and 0.06 into Yorkey’s Knob [9]. Alternatively, the failure to reach fixation in these areas could be due to a combination of migration and infection loss, which would mean that migration rates of uninfected, gravid females may be lower than 0.06.

## Conclusions

Our study has provided empirical evidence for three processes predicted to slow down the spread of *w*Mel in *Ae. aegypti*, using a landscape genomics analytical framework and molecular assays of *Wolbachia* infection. This approach could be extended to other host/*Wolbachia* systems that are increasingly considered for the biocontrol of disease vectors and pests. Non-perfect maternal transmission of *w*Mel in *Ae. aegypti* may not occur in other *Wolbachia* strains such as *w*AlbB, whose density shows greater constancy under fluctuating high temperatures [37], Also, it is as yet unclear whether the observed transmission failure of *w*Mel occurs at a high enough frequency to affect invasion dynamics, and a more extensive field test of transmission fidelity will be necessary to derive an accurate estimate of *μ*. On the other hand, regardless of the *Wolbachia* strain deployed, presence of barriers like highways and leptokurtic dispersal are potential challenges for any *Wolbachia* invasion strategy requiring spatial spread [19].

## Acknowledgements

We would like to thank Eliminate Dengue Cairns, particularly field officer Angela Caird, for assistance with ovitrap deployment. We thank the Cairns householders participating in the study for granting permission to deploy ovitraps on their property, as well as Scott Ritchie and Christopher Paton from the Centre for Biosecurity in Tropical Infectious Diseases, James Cook University, for assisting with the processing of field collections. The National Health and Medical Research Council provided funding for this research through a Program grant and Fellowship grant to A.A. Hoffmann.

## Data Archiving

Demultiplexed fastq files have been deposited at NCBI SRA under [name].

## Author Contributions

A.A. Hoffmann, G. Rašić, and T. L. Schmidt conceived of and designed the study.

T. L. Schmidt, G. Rašić, and I. Filipović collected and dried the samples.

T. L. Schmidt performed the laboratory work and conducted the analyses, with assistance from G. Rašić, and computational support from I. Filipović.

T. L. Schmidt wrote the manuscript with assistance from A.A. Hoffmann and G. Rašić.

